# Ligand Gaussian accelerated Molecular Dynamics 3 (LiGaMD3): Improved Calculations of Binding Thermodynamics and Kinetics of Both Small Molecules and Flexible Peptides

**DOI:** 10.1101/2024.05.06.592668

**Authors:** Jinan Wang, Yinglong Miao

## Abstract

Binding thermodynamics and kinetics play critical roles in drug design. However, it has proven challenging to efficiently predict ligand binding thermodynamics and kinetics of small molecules and flexible peptides using conventional Molecular Dynamics (cMD), due to limited simulation timescales. Based on our previously developed Ligand Gaussian accelerated Molecular Dynamics (LiGaMD) method, we present a new approach, termed “LiGaMD3”, in which we introduce triple boosts into three individual energy terms that play important roles in small-molecule/peptide dissociation, rebinding and system conformational changes to improve the sampling efficiency of small-molecule/peptide interactions with target proteins. To validate the performance of LiGaMD3, MDM2 bound by a small molecule (Nutlin 3) and two highly flexible peptides (PMI and P53) were chosen as model systems. LiGaMD3 could efficiently capture repetitive small-molecule/peptide dissociation and binding events within 2 microsecond simulations. The predicted binding kinetic constant rates and free energies from LiGaMD3 agreed with available experimental values and previous simulation results. Therefore, LiGaMD3 provides a more general and efficient approach to capture dissociation and binding of both small-molecule ligand and flexible peptides, allowing for accurate prediction of their binding thermodynamics and kinetics.

## Introduction

Both small molecules and peptides are important sources of novel drugs targeting many important biological processes^1^. It is critical to understand the binding mechanisms of small molecules and peptides to their target proteins, which not only deepens our understanding of fundamental biological processes but also facilitates the development of more potent and selective drugs for treating human diseases^2^. Several experimental techniques^3, 4^ have been developed to explore the binding interactions between the protein and small molecule or peptide. For example, structural biology techniques including X-ray crystallography and cryo-electron microscopy (cryo-EM) have been widely used to determine the complex structures of protein-small molecule and protein-peptide complexes^4^. Recently, significant advancements in Deep Learning methodologies such as AlphaFold^5^ and RoseTTAFold All-Atom (RFAA)^6^ have led to accurate prediction of protein-small molecule or protein-peptide complex structures. However, such techniques provide only static snapshots of protein-small molecule or protein-peptide interactions. It is still challenging to capture small-molecule/peptide binding and dissociation processes and determine potential intermediate states of small-molecule/peptide binding to their target proteins, which are also important for drug design^7^.

Recently, drug binding kinetics has been recognized to be valuable for drug design^8, 9^. The drug dissociation rate (*k*_*off*_) appears to correlate with drug efficacy better than the binding free energy^8, 9^. However, drug binding kinetic rates have proven more challenging to predict, due to the slow processes of drug dissociation and binding^9, 10^.With remarkable advancements in computer hardware and methodological developments, conventional Molecular Dynamics (cMD) simulations are now able to capture spontaneous small-molecule/peptide binding to their target proteins and predict corresponding association rates (k_on_)^11, 12^. However, it remains a difficult challenge in applying cMD to capture repetitive small-molecule/peptide dissociation and rebinding processes within accessible timescales, thereby hindering the accurate prediction of small-molecule/peptide binding kinetic rates^13^. Shan et al.^13^ successfully captured spontaneous binding of the Dasatinib drug to its target Src kinase and accurately predicted the ligand association rate based on tens-of-microsecond cMD simulations. Tens-of-microsecond cMD simulations^12^ have successfully captured repetitive binding and dissociation events of six small-molecule fragments with very weak millimolar binding affinities to the protein FKBP, allowing accurate prediction of fragment binding free energies. However, no dissociation events have been observed for typical ligand molecules in the cMD simulations. Furthermore, capturing peptide dissociation and rebinding processes poses an even more challenging task for cMD, given that peptides are known to induce significant conformational changes upon binding^14^ and the timescales for the peptide dissociation are even longer^15^. For example, cMD simulations with elevated temperature conducted for 200 μs using the Anton specialized supercomputer have captured 70 binding and unbinding events between an intrinsically disordered protein fragment of the measles virus nucleoprotein and the X domain of the measles virus phosphoprotein complex, shedding light on the detailed understanding of the peptide’s “folding-upon-binding” mechanism^16^. Despite these advancements, it is still rather challenging for cMD to effectively simulate binding and dissociation of typical small molecule or peptide to their target proteins.

Enhanced sampling methods^17^ have been developed to extend the accessible timescales of MD simulations. These methods include Metadynamics^18^, Steered MD^19, 20^, Umbrella Sampling^19, 21^, Replica Exchange MD ^22, 23^, Random Acceleration Molecular Dynamics (RAMD) ^24^, Scaled MD ^25^, accelerated MD (aMD) ^26^, Gaussian accelerated MD (GaMD) ^27, 28^, Markov State Model (MSM)^29, 30^, Weighted Ensemble^31, 32^, and so on. Metadynamics^14, 33^ simulations utilizing carefully chosen collective variables (CVs) have successfully predicted the peptide binding and dissociation rates in the systems. Particular, 27 μs Metadynamics simulations of the peptide P53 binding to the MDM2^33^ predicted values of (k_on_, k_off_) at (0.43±0.22x10^7^M^-1^s^-1^, 0.7±0.4s^-1^), showing good agreement with the corresponding experimental values of (0.92x10^7^M^-1^s^-1^, 2.06s^-1^). Similarly, Weighted Ensemble^34^ of a total amount of *∼*120 μs cMD simulations with implicit solvent model for the P53-MDM2 system predicted a highly consistent binding kinetic rate (*k*_*on*_) of 7 s^-1^. MSM^35^ analysis based on a total of 831 μs cMD simulations for peptide P53 binding to MDM2 accurately predicted values of k_on_ and k_off_ at 0.019x10^7^ M^-1^s^-1^ and 2.5 s^-1^, respectively. Another MSM built on hundreds-of-microsecond cMD and Hamiltonian replica exchange simulations has been implemented to characterize binding and dissociation of the PMI peptide to the MDM2^36^. The predicted values of (k_on_, k_off_) were (300x10^7^M^-1^s^-1^, 0.125-1.13s^-1^), being comparable to the corresponding experimental values of (52.7x10^7^M^-1^s^-1^, 0.037s^-1^). Nevertheless, MSM and Weighted Ensemble require expensive and exceedingly long simulations. GaMD was developed to provide both unconstrained enhanced sampling and free energy calculations of large biomolecules^27, 28^. It works by applying a harmonic boost potential to reduce system energy barriers. The boost potential exhibits a near Gaussian distribution, which enables accurate reweighting of the free energy profiles through cumulant expansion to the second order^27, 28^. Recently, novel Ligand GaMD (LiGaMD)^37^ and LiGaMD2^38^ approaches have been developed to more efficiently sample small-molecule dissociation and rebinding processes, offering accurate prediction of ligand binding thermodynamics and kinetics. In LiGaMD, a selective boost is specifically applied at the ligand’s non-bonded interaction potential energy^39^. In LiGaMD2, the selective boost extended to the essential potential energy of both the ligand and surrounding residues in the protein pocket, which significantly improved the sampling of ligand dissociation and rebinding in a closed binding pocket^38^. An increasing amount of studies^23, 30, 40^ demonstrated the pivotal role of non-bonded interaction potentials in ligand binding, along with the crucial structural flexibility of proteins and peptides^30, 41, 42, 43^. Building upon the successes of LiGaMD and LiGaMD2, which primarily focused on small-molecules, here we introduce a more general approach, termed LiGaMD3, for binding simulations of both small-molecules and flexible peptides. In LiGaMD3, three distinct boosts are applied: one on the non-bonded interaction energy of the substrate, the second one on the remaining non-bonded potential energy of the system, and a third one on the system bonded potential energy. These boosts are designed to accelerate the substrate dissociation, facilitate substrate rebinding, and promote the system conformational changes, respectively. MDM2^44^, a well-known oncology protein involved in regulating diverse cellular signaling pathways, serves as an ideal model system for investigating the binding and dissociation of both small molecules and highly flexible peptides. Notably, this system has been extensively explored through experimental studies and simulations as mentioned above, highlighting its critical role in drug discovery. Therefore, MDM2 bound by small-molecule drugs and peptides were chosen as model systems in this study. Through two microsecond LiGaMD3 simulations, we successfully captured repetitive ligand and peptide binding and dissociation processes across all MDM2 systems. LiGaMD3 facilitated highly efficient and accurate predictions of ligand and peptide binding thermodynamics and kinetics, being consistent with experimental binding free energies and kinetic rates.

## Methods

### LiGaMD3: Triple boost for ligand dissociation and rebinding

We consider a system of small-molecule/peptide *L* binding to a protein *P* in a biological environment *E*. The system comprises of *N* atoms with their coordinates 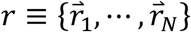 and momenta 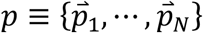. The system Hamiltonian can be expressed as:

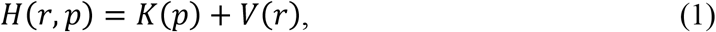

where *K* (*p*) and *V* (*r*) are the system kinetic and total potential energies, respectively. We decompose the potential energy into the following terms:

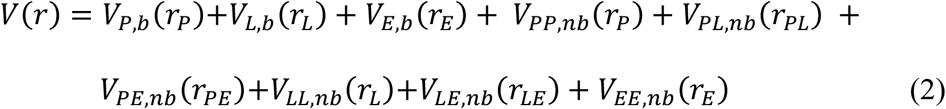

where *V*_*P,b*_, *V*_*L,b*_ and *V*_*E,b*_ are the bonded potential energies of protein P, small-molecule/peptide *L* and environment *E*, respectively. *V*_*P,nb*_, *V*_*L,nb*_ and *V*_*E,nb*_ are the self non-bonded potential energies in the protein P, small-molecule/peptide *L* and environment *E*, respectively. *V*_*PL,nb*_, *V*_*PE,nb*_ and *V*_*LE,nb*_ are the non-bonded interaction energies between P-L, P-E and L-E, respectively. According to classical molecular mechanics force fields^45^, the non-bonded potential energies are usually calculated as:

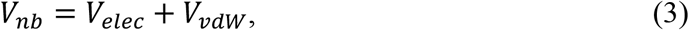

where *V*_*elex*_ and *V*_*vdw*_ are the system electrostatic and van der Waals potential energies. The bonded potential energies are usually calculated as

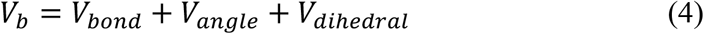

where *V*_*bond*_, *V*_*angle*_ and *V*_*dihedral*_ are the system bond, angle and dihedral potential energies. In LiGaMD3, the essential non-bonded interaction potential energy of the ligand is defined as:

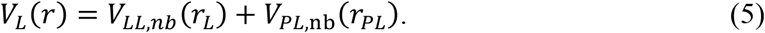

We add a boost potential selectively to the *V*_*L*_ (*r*) according to the GaMD algorithm:

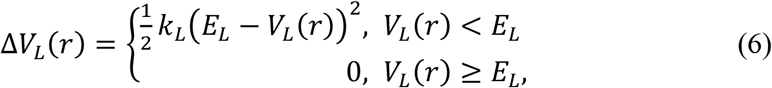

where *E*_*L*_ is the threshold energy for applying boost potential and *k*_*L*_ is the harmonic constant. The LiGaMD3 simulation parameters are derived similarly as in the previous GaMD^46^, LiGaMD^47^, and LiGaMD2^38^. When *E* is set to the lower bound as the system maximum potential energy (*E=V*_*max*_), the effective harmonic force constant *k*_0_ can be calculated as:

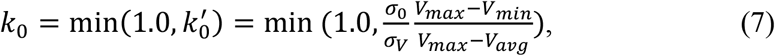

where *V*_*max*_, *V*_*min*_, *V*_*avg*_ and *σ*_*V*_ are the maximum, minimum, average and standard deviation of the boosted system potential energy, and *σ*_0_ is the user-specified upper limit of the standard deviation of Δ*V* (e.g., 10*k*_*B*_T) for proper reweighting. The harmonic constant is calculated as *k* = *k*_0_ 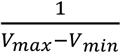 with 0 < *k*_0_ ≤ 1. Alternatively, when the threshold energy *E* is set to its upper bound 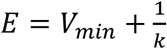, *k*_0_ is set to:

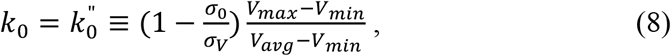

if 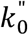 is found to be between *0* and *1*. Otherwise, *k*_0_ is calculated using Eqn. (7).

In addition to selectively boosting the essential non-bonded interaction potential energy of the ligand, another boost potential is applied on the remaining non-bonded potential energy of the system (*V*_*D*_ (*r*)) to facilitate ligand rebinding:

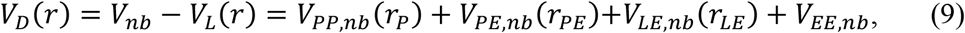

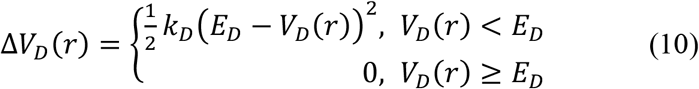

where *V*_*D*_ is the total system potential energy other than the essential non-bonded ligand potential energy ligand, *E*_*D*_ is the corresponding threshold energy for applying the second boost potential and *k*_*D*_ is the harmonic constant.

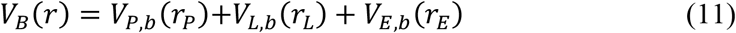

The third boost potential is calculated using the total bonded potential energy of the system as:

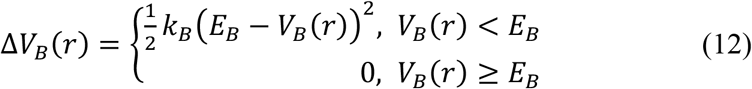

This leads to LiGaMD3 with a triple-boost potential Δ*V*(*r*) = Δ*V*_*L*_(*r*) + Δ*V*_*D*_(*r*) + Δ*V*_*B*_ (*r*).

### Energetic Reweighting of LiGaMD3

To calculate potential of mean force (PMF)^48^ from LiGaMD3 simulations, the probability distribution along a reaction coordinate is written as *p*^*^(*A*). Given the boost potential Δ*V*(*r*) of each frame, *p*^*^(*A*) can be reweighted to recover the canonical ensemble distribution, *p* (*A*), as:

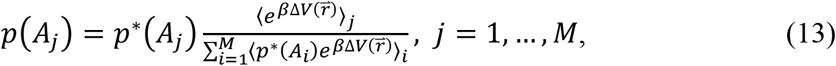

where *M* is the number of bins, *β* = *k*_*B*_*T* and 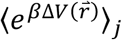 is the ensemble-averaged Boltzmann factor of 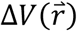 for simulation frames found in the *j*^th^ bin. The ensemble-averaged reweighting factor can be approximated using cumulant expansion:

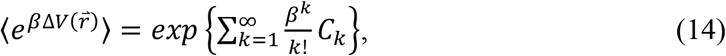

where the first two cumulants are given by

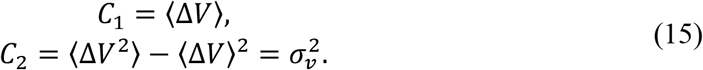

The boost potential obtained from LiGaMD3 simulations usually follows near-Gaussian distribution. Cumulant expansion to the second order thus provides a good approximation for computing the reweighting factor^28, 49^. The reweighted free energy *F* (*A*) = −*k*_*B*_*T* ln *p* (*A*) is calculated as:

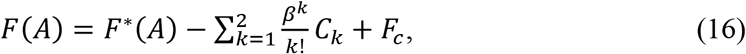

where *F*^*^(*A*) = −*k*_*B*_*T* ln *p*^*^(*A*) is the modified free energy obtained from LiGaMD2 simulation and *F*_*e*_ is a constant.

### Ligand binding kinetics obtained from reweighting of LiGaMD3 Simulations

Reweighting of ligand binding kinetics from LiGaMD3 simulations followed a similar protocol using Kramers’ rate theory that has been recently implemented in kinetics reweighting of the GaMD^50, 51, 52^. Provided sufficient sampling of repetitive ligand dissociation and binding in the simulations, we record the time periods and calculate their averages for the ligand found in the bound (*τ*_*B*_) and unbound (*τ*_*U*_) states from the simulation trajectories. The *τ*_*B*_ corresponds to residence time in drug design^53^. Then the ligand dissociation and binding rate constants (*k*_*off*_ and *k*_*on*_) were calculated as:

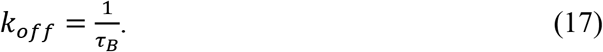

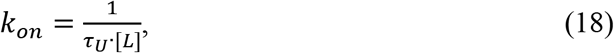

where [L] is the ligand concentration in the simulation system.

According to Kramers’ rate theory, the rate of a chemical reaction in the large viscosity limit is calculated as^52^:

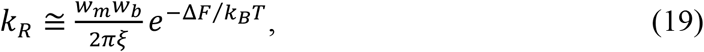

where *w*_*m*_ and *w*_*b*_ are frequencies of the approximated harmonic oscillators (also referred to as curvatures of free energy surface^54^) near the energy minimum and barrier, respectively, *ξ* is the frictional rate constant and Δ*F* is the free energy barrier of transition. The friction constant *ξ* is related to the diffusion coefficient *D* with *ξ* = *k*_*B*_*T*/*D*. The apparent diffusion coefficient *D* can be obtained by dividing the kinetic rate calculated directly using the transition time series collected directly from simulations by that using the probability density solution of the Smoluchowski equation^55^. In order to reweight ligand kinetics from the LiGaMD3 simulations using the Kramers’ rate theory, the free energy barriers of ligand binding and dissociation are calculated from the original (reweighted, ***ΔF***) and modified (no reweighting, ***ΔF\****) PMF profiles, similarly for curvatures of the reweighed (*w*) and modified (*w*^*^, no reweighting) PMF profiles near the ligand bound (“B”) and unbound (“U”) low-energy wells and the energy barrier (“Br”), and the ratio of apparent diffusion coefficients from simulations without reweighting (modified, *D*^*^) and with reweighting (*D*). The resulting numbers are then plugged into Eq. (17) to estimate accelerations of the ligand binding and dissociation rates during LiGaMD3 simulations^52^, which allows us to recover the original kinetic rate constants.

### System Setup

The complex structures of MDM2 bound by the Nutlin 3 drug, P53 and PMI were obtained from the 5C5A, 1YCR and 3EQS PDB files, respectively. The AMBER ff14SB force field^56^ was used for the protein and peptide. The GAFF2 force field^57^ with AM1-BCC charge was used for the Nutlin 3 small molecule. Each system was solvated in a periodic box of TIP3P water molecules with a distance of 18 Å from the solute to the box edge using *tleap*. Therefore, the ligand/peptide concentration was 0.00335 M in the simulation system. Appropriate number of Na+/Cl-ions were added to achieve system neutrality.

### Simulation Protocol

Each system was energy minimized and gradually heated to 300 K in 1 ns with the Langevin thermostat and harmonic restraints of 1.0 kcal/mol/Å^2^ on all non-hydrogen atoms of the protein and the ligand using the AMBER23 software^58^. The simulation system was firstly energy minimized with 1.0 kcal/mol/Å^2^ constraints on the heavy atoms of the proteins, including the steepest descent minimization for 50,000 steps and conjugate gradient minimization for 50,000 steps. The system was then heated from 0 K to 300 K for 200 ps. It was further equilibrated using the NVT ensemble at 300 for 200 ps and the NPT ensemble at 300 K and 1 bar for 1 ns with 1 kcal/mol/Å^2^ constraints on the heavy atoms of the protein, followed by 2 ns short cMD without any constraint. The LiGaMD3 simulations proceeded with 2 ns short cMD to collect the potential statistics, 50.0 ns LiGaMD3 equilibration after adding the boost potential and then three independent 2,000 ns production runs. It provided more powerful sampling to set the threshold energy for applying the boost potential to the upper bound (i.e., *E* = *V*_min_+1/*k*) in our previous study ligand dissociation and binding using LiGaMD^51^. Therefore, the threshold energy for applying the ligand essential non-bonded potential (first boost) and the remaining non-bonded potential energy of the system (second boost) were set to the upper bound in the LiGaMD3 simulations. The threshold energy for applying the third boost to the system bonded energy potential was set to the lower bound. In order to observe ligand/peptide dissociation during LiGaMD3 production simulations while keeping the boost potential as low as possible for accurate energetic reweighting, the (*σ*_0P_, *σ*_0D,_ *σ*_0B_) parameters were set to (2.0 kcal/mol, 6.0 kcal/mol, 6.0 kcal/mol) for the LiGaMD3 simulations of the MDM2 bound by the Nutlin 3, PMI and P53. LiGaMD3 production simulation frames were saved every 0.4 ps for analysis. In the LiGaMD simulations performed for comparison, the (*σ*_0P_, *σ*_0D_) parameters were set to (4.8 kcal/mol, 6.0 kcal/mol) for simulations of the MDM2-Nutlin 3 system and (7.0 kcal/mol, 6.0 kcal/mol) for MDM2-PMI system, respectively. The threshold energies for applying the boosts in the LiGaMD simulations were set to the upper bound. Example simulation files of the LiGaMD3 simulations of the MDM2-Nutlin 3 system are included in the Supporting Information.

### Simulation Analysis

The VMD^59^ and CPPTRAJ^60^ tools were used for simulation analysis. The number of ligand dissociation and binding events (*ND* and *NB*) and the ligand/peptide binding and unbinding time periods (*τB* and *τU*) were recorded from individual simulations (**Tables 1 & S1**). With high fluctuations, *τB* and *τU* were recorded for only the time periods longer than 1 ns. The 1D and 2D free energy profiles, as well as the ligand binding free energy, were calculated through energetic reweighting of the LiGaMD3 simulations. The center-of-mass distance between the small-molecule/peptide and the protein pocket (defined by protein residues within 5 Å of the ligand, denoted as d_MDM2-substrate_) and small-molecule heavy atom or peptide backbone RMSDs relative to the PDB structures with the protein aligned were chosen as the reaction coordinates. The bin size was set to 1.0 Å. The cutoff for the number of simulation frames in one bin was set to 500. The ligand binding free energies (*ΔG*) were calculated using the binding kinetic rates as Δ*G* = −RTLn*k*_*off*_/*k*_o*n*_. The ligand dissociation and binding rate constants (*k*_*on*_ and *k*_*off*_) were calculated from the LiGaMD3 and LiGaMD simulations with their accelerations analyzed using the Kramers’ rate theory (**Table S2**).

**Table 1.** Summary of LiGaMD3 simulations performed on small molecule and peptide binding to the MDM2. *ΔV* is the total boost potential. *N*_*D*_ and *N*_*B*_ are the number of observed ligand dissociation and binding events, respectively. *ΔG*_*sim*_ and *ΔG*_*exp*_ are the ligand-MDM2 binding free energies obtained from LiGaMD3 simulations and experiments, respectively. ^a^ The simulation binding free energy is estimated using *ΔG*_*sim*_=-*RT* Ln(*k*_*off*_/*k*_*on*_).

| System | Method | ID | N<sub>B</sub> | N<sub>D</sub> | $\Delta V$<br>(kcal/mol) | $\Delta G_{sim}^a$<br>(kcal/mol) | $\Delta G_{exp}$<br>(kcal/mol) |
| --- | --- | --- | --- | --- | --- | --- | --- |
| MDM2-Nutline3 | LiGaMD | Sim1 | 5 | 6 | 94.92±4.01 | -10.18±2.22 | -10.96 |
|  |  | Sim2 | 3 | 3 |  |  |  |
|  |  | Sim3 | 3 | 4 |  |  |  |
| MDM2-Nutline3 | LiGaMD3 | Sim1 | 6 | 5 | 12.94±4.05 | -11.02±0.59 |  |
|  |  | Sim2 | 7 | 7 |  |  |  |
|  |  | Sim3 | 5 | 5 |  |  |  |
| MDM2-PMI | LiGaMD3 | Sim1 | 4 | 4 | 45.11±7.04 | -11.86±1.16 | -12.02 |
|  |  | Sim2 | 6 | 5 |  |  |  |
|  |  | Sim3 | 6 | 6 |  |  |  |
| MDM2-P53 | LiGaMD3 | Sim1 | 5 | 6 | 45.59±7.03 | -10.59±0.11 | -9.27 |
|  |  | Sim2 | 4 | 5 |  |  |  |
|  |  | Sim3 | 5 | 6 |  |  |  |

## Results

### Microsecond LiGaMD3 simulations captured repetitive small-molecule and peptide dissociation and rebinding to the MDM2

Both LiGaMD and LiGaMD3 have effectively captured the binding and dissociation processes of the Nutlin 3 small molecule to the MDM2 protein across all three independent 2,000 ns simulations (**Figs. 1B & 1C**). However, LiGaMD encountered difficulty in capturing the rebinding of the PMI peptide to the protein, as no frames with peptide RMSD < 5 Å were observed in all three 2,000 ns simulations (**Fig. 1F**). In contrast, LiGaMD3 demonstrated consistent performance in successfully capturing the repetitive binding and dissociation of the PMI peptide in the MDM2 (**Fig. 1E**). Moreover, an additional system wherein MDM2 is bound by the peptide P53 was included to further evaluate the performance of LiGaMD3 (**Fig. 2**). LiGaMD3 could capture multiple times of P53 binding and dissociation in 2000ns simulations (**Fig. 2**).

**Figure 1.**
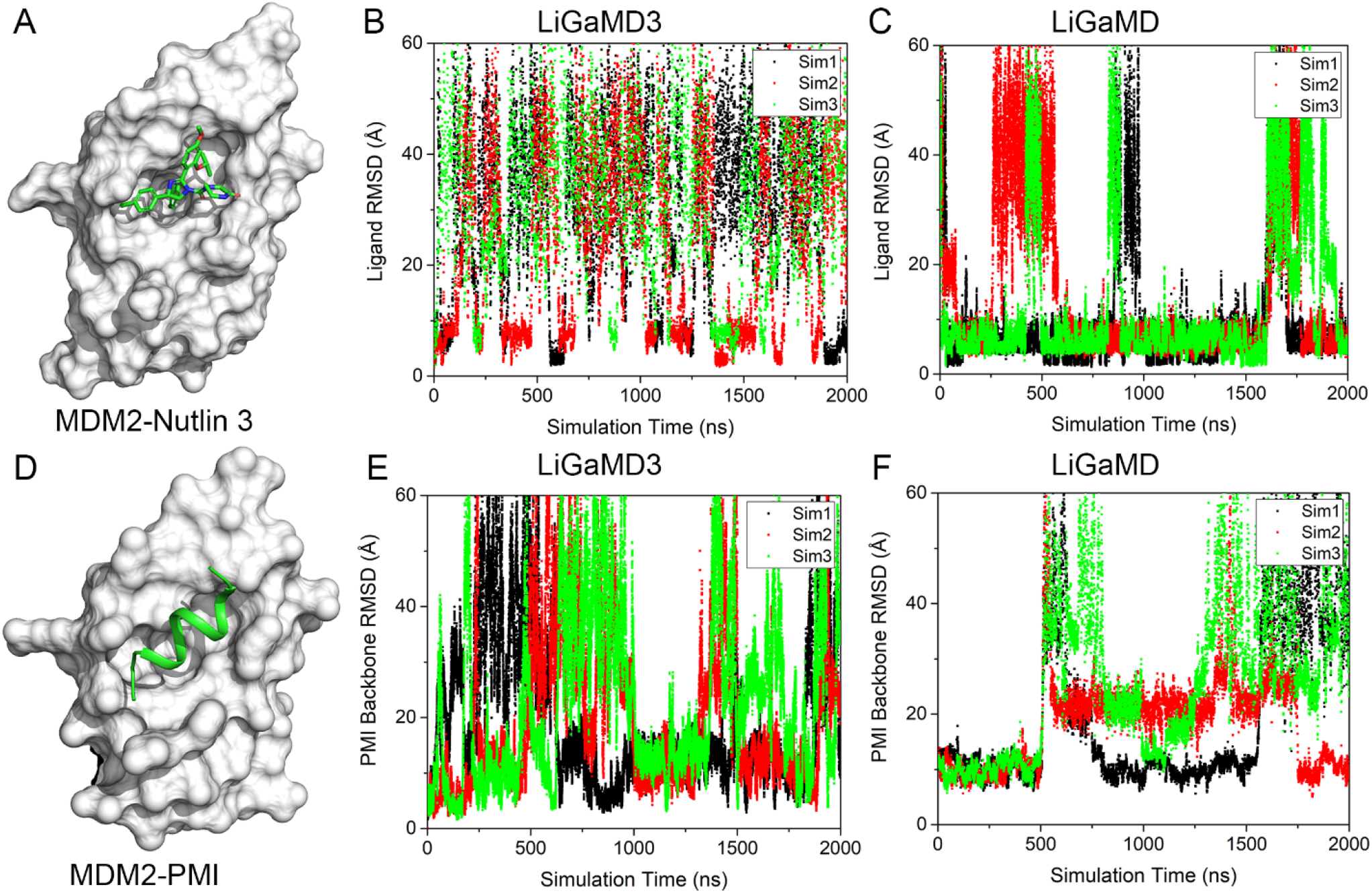
Comparison of LiGaMD and LiGaMD3 simulations on the MDM2 protein bound by the Nutlin 3 small molecule and PMI peptide: Computational models of the MDM2 bound by the Nutlin 3 small molecule (**A**) and PMI peptide (**D**); Time courses of ligand root-mean-square deviation (RMSD) relative to the experimental bound structure (PDB ID: 5C5A) in the MDM2-Nutlin 3 system calculated from LiGaMD3 (**B**) and LiGaMD (**C**) simulations, respectively. Time courses of peptide RMSD relative to the experimental bound structure (PDB ID: 3EQB) in the MDM2-PMI calculated from LiGaMD3 (**E**) and LiGaMD2 (**F**) simulations, respectively.

**Figure 2.**
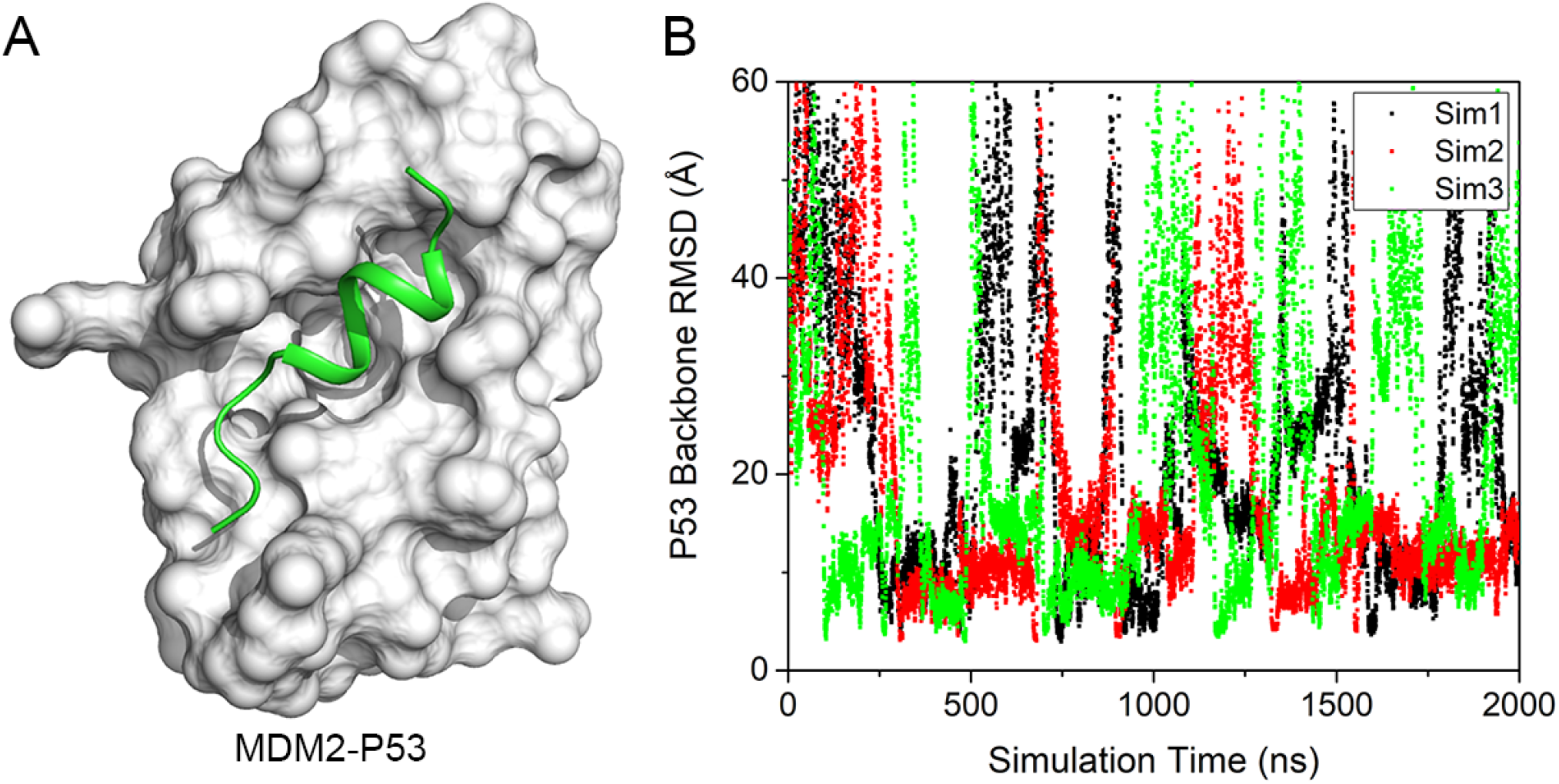
(**A**) Computational model of the MDM2 bound by the P53 peptide; (**B**) Time courses of peptide P53 RMSD relative to the experimental bound structure (PDB ID: 1YCR) in the MDM2-P53 system calculated from LiGaMD3 simulations.

The LiGaMD3 simulations of the MDM2-Nutlin 3 system yielded an average boost potential of 12.92 kcal/mol with a standard deviation of 4.05 kcal/mol (**Table 1**). In contrast, achieving ligand dissociation and binding required a significantly larger boost in LiGaMD, with an average boost potential of 94.92 kcal/mol and a standard deviation of 4.01 kcal/mol (**Table 1**). For the MDM2-PMI and MDM2-P53 systems, LiGaMD3 simulations recorded average boost potentials of 45.11±7.04 kcal/mol and 45.59±7.03 kcal/mol, respectively (**Table 1**). Notably, the boost potentials applied in simulations of peptide-protein systems are much higher compared to that of small molecule-MDM2 system. LiGaMD3 required substantially smaller boosts than LiGaMD simulations, indicating its enhanced efficiency in sampling, which was also advantageous for accurate energetic and kinetic reweighting.

RMSDs of the small-molecule/peptide relative to their experimental bound structures with the MDM2 protein aligned were computed (**Figs. 1**, 2 & **Table 1**) to calculate the number of small-molecule/peptide dissociation (*N*_*D*_) and binding (*N*_*B*_) events in each of the 2,000 ns LiGaMD3 simulations. With close examination of the ligand/peptide binding trajectories, RMSD cutoffs of the ligand unbound and bound states were set to >15 Å and <5.0 Å, respectively. Due to fluctuations in small-molecule/peptide-protein interactions, we recorded only the corresponding binding and dissociation events that lasted for more than 1.0 ns. In 2,000 ns simulations of the MDM2-Nutlin 3 system, LiGaMD3 consistently captured 5-7 binding and 5-7 dissociation events, whereas LiGaMD captured only 3-5 binding and 3-6 dissociation events (**Fig. 1 & Table 1**). The total number of binding events recorded in LiGaMD3 was 18, compared to 11 in LiGaMD. The total number of dissociation events in LiGaMD3 and LiGaMD was 17 and 13, respectively. Hence, LiGaMD3 demonstrated improved efficiency in capturing both binding and dissociation events compared to LiGaMD. Additionally, no rebinding events were observed in LiGaMD simulations of PMI to MDM2 (**Fig. 1F**), whereas each 2,000 ns LiGaMD3 simulation successfully captured 4-6 binding and 4-6 dissociation events, indicating its superior capability in capturing flexible peptide-protein interactions (**Fig. 1E & Table 1**). Similar numbers of peptide dissociation (5-6) and binding (4-5) events were observed in simulations of the MDM2-P53 system (**Fig. 2 & Table 1**). In summary, LiGaMD3 simulations successfully captured repetitive dissociation and rebinding events of both small-molecules and flexible peptides to the MDM2 on three model systems: Nutlin 3 bound to MDM2 (MDM2-Nutlin 3), PMI bound to MDM2 (MDM2-PMI), and P53 bound to MDM2 (MDM2-P53) (**Figs. 1, 2 & S1**).

### Ligand and peptide binding kinetic rates and free energies calculated from LiGaMD3 agreed well with experimental data

The successful simulations of repetitive small-molecule and flexible peptide binding and dissociation in LiGaMD3 allowed us to predict the small-molecule and peptide binding kinetic rate constants (**Fig. S2 & Table 2**). We recorded the time periods for the small-molecule and peptide found in the bound (*τ*_*B*_) and unbound (*τ*_*U*_) states throughout the LiGaMD and LiGaMD3 simulations. Without reweighting, the binding rate constant (*k*_*on*_*\**) and dissociation rate constant (*k*_*off*_*\**) for Nutlin 3 were directly calculated from the LiGaMD trajectories as 1.69 ± 0.28×10^9^ M^-1^.s^-1^ and 3.68 ± 2.10×10^6^ s^-1^ (**Table 2**). In comparison, in LiGaMD3, these rate constants were calculated as 1.16 ± 0.28×10^9^ M^-1^.s^-1^ and 2.77 ± 0.75×10^7^ s^-1^, respectively (**Table 2**). The peptide binding rate constants (*k*_*on*_*\**) were directly calculated from the LiGaMD3 trajectories as 8.57 ± 1.37×10^8^ M^-1^.s^-1^ and 1.02±0.29 × 10^9^ M^-1^.s^-1^ for the MDM2-PMI and MDM2-P53 systems, respectively (**Table 2**).

**Table 2.** Comparison of kinetic rates obtained from experimental data and LiGaMD3 simulations for ligand binding to MDM2. *k*_*on*_ and *k*_*off*_ are the kinetic dissociation and binding rate constants, respectively, from experimental data or LiGaMD3 simulations with reweighting using Kramers’ rate theory. 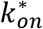 and 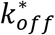 are the accelerated kinetic dissociation and binding rate constants calculated directly from LiGaMD3 simulations without reweighting.

| System | Method | $k_{on} (M^{-1} \cdot s^{-1})$ | $\Delta \log(k_{on})$ | $k_{off} (s^{-1})$ | $\Delta \log(k_{off})$ | $k_{on}^* (M^{-1} \cdot s^{-1})$ | $k_{off}^* (s^{-1})$ |
| --- | --- | --- | --- | --- | --- | --- | --- |
| MDM2-Nutlin | Experiment | $3.3 \times 10^7$ | - | 0.48 | - | - | - |
| | LiGaMD | $5.26 \pm 0.65 \times 10^9$ | 1.82 | $69.51 \pm 58.37$ | 1.36 | $1.69 \pm 0.28 \times 10^9$ | $3.68 \pm 2.10 \times 10^6$ |
| | LiGaMD3 | $8.29 \pm 4.80 \times 10^8$ | 1.37 | $15.45 \pm 4.69$ | 1.39 | $1.16 \pm 0.31 \times 10^9$ | $2.77 \pm 0.75 \times 10^7$ |
| MDM2-PMI | Experiment | $5.27 \times 10^8$ | - | 0.037 | - | - | - |
| | LiGaMD3 | $5.76 \pm 4.80 \times 10^8$ | 0.04 | $2.66 \pm 1.73$ | 1.85 | $8.51 \pm 1.37 \times 10^8$ | $3.41 \pm 0.96 \times 10^7$ |
| MDM2-P53 | Experiment | $9.2 \times 10^6$ | - | 2.06 | - | - | - |
| | LiGaMD3 | $8.03 \pm 7.42 \times 10^8$ | 1.94 | $28.0 \pm 19.2$ | 1.13 | $1.02 \pm 0.29 \times 10^9$ | $1.95 \pm 0.73 \times 10^7$ |

Next, we performed reweighting on the LiGaMD and LiGaMD3 simulations of ligand-MDM2 systems to calculate acceleration factors for the small-molecule/peptide binding and dissociation processes (**Table S2**) and to recover the original kinetic rate constants using the Kramers’ rate theory (**Table 2**). In the LiGaMD simulations, the dissociation free energy barrier (*ΔF*_*off*_) significantly decreased from 9.10±1.26 kcal/mol in the reweighted PMF profiles to 2.77±1.76 kcal/mol in the modified PMF profiles for the system of MDM2-Nutlin 3 (**Fig. S1** and **Table S2**). Similarly, for the MDM2-Nutlin 3, MDM2-PMI, and MDM2-P53 systems, the dissociation free energy barrier (*ΔF*_*off*_) significantly decreased from 9.65±0.76, 9.05±0.23, 7.23±0.40 kcal/mol in the reweighted PMF profiles to 0.79±0.10, 0.89±0.07, 0.67±0.18 kcal/mol in the modified PMF profiles in LiGaMD3 simulations, respectively (**Table S1 and Fig. S1**). Curvatures of the reweighed (*w*) and modified (*w*^*^, no reweighting) free energy profiles were calculated near the ligand Bound (“B”) and Unbound (“U”) low-energy wells and the energy barrier (“Br”), as well as the ratio of apparent diffusion coefficients calculated from LiGaMD and LiGaMD3 simulations with reweighting (*D*) and without reweighting (modified, *D*^*^) (**Table S2**). According to the Kramers’ rate theory, the association and dissociation of the Nutline 3 small molecule in LiGaMD were accelerated by 0.32 and 5.30×10^4^ times. In contrast, in LiGaMD3, the association and dissociation of the Nutlin 3 were accelerated by 1.40 and 1.79×10^6^ times, respectively. Moreover, the association of the peptide in the LiGaMD3 was accelerated by 1.47 and 1.27 times for the MDM2-PMI and MDM2-P53 systems, respectively. While the peptide dissociation was significantly accelerated by 1.28×10^7^ and 6.96×10^5^ times for the MDM2-PMI and MDM2-P53 systems, respectively. Therefore, the reweighted *k*_*on*_ in the MDM2-Nutlin 3 system with LiGaMD and LiGaMD3 were calculated as 5.26±0.65×10^9^ M^-1^.s^-1^ and 8.29±4.80×10^8^ M^-1^.s^-1^, respectively, being in consistent with the experimental values of 3.3×10^7^ M^-1^.s^-1^. Similarly, for the MDM2-PMI and MDM2-P53 systems, the reweighted *k*_*on*_ values were predicted as 5.76±4.80×10^8^ and 8.03±7.42×10^8^, respectively, being consistent with the corresponding experimental values^61^ of 5.27×10^8^ and 9.20×10^6^ M^-1^.s^-1^(**Table 2**). The reweighted *k*_*off*_ values for the Nutlin 3 in the MDM2-Nutline 3 with LiGaMD and LiGaMD3 were calculated as 69.51±58.37 s^-1^ and 15.45±4.69 s^-1^, being in accordance with the experimental of 0.48 s^-1^. For the peptide in the MDM2-PMI and MDM2-P53 systems, the reweighted peptide *k*_*off*_ were calculated from LiGaMD3 simulations as 2.66±1.73, 28.0±19.2 s^-1^, in agreement with the corresponding experimental values^61^ of 0.037 and 2.06 s^-1^, respectively.

Based on the ligand binding kinetic rates (*k*_*on*_ and *k*_*off*_), we calculated the ligand binding free energies as Δ*G* = −RTLn*k*_*off*_/*k*_*on*_. The resulting binding free energies in the MDM2-Nutlin 3 system with LiGaMD and LiGaMD3 were -10.18±2.22 kcal/mol and -11.02±0.59 kcal/mol, respectively, demonstrating high consistency with the experimental value of -10.96 kcal/mol. In the MDM2-PMI and MDM2-P53 systems (**Table 1**), the calculated peptide binding free energy values were -11.86±1.16 kcal/mol and -10.59±0.11 kcal/mol, exhibiting strong agreement with the corresponding experimental values of -12.02 kcal/mol and -9.27 kcal/mol respectively. The root-mean square error (RMSE) of binding free energy for the three systems was only 0.94 kcal/mol. Hence, LiGaMD3 simulations achieved both efficient sampling and accurate small-molecule/peptide binding thermodynamics and kinetics calculations.

### Multiple ligand binding and dissociation pathways were identified from LiGaMD3 simulations

We closely examined the LiGaMD3 trajectories to explore the pathways involved in the small-molecule/peptide binding and dissociation of the MDM2. Two primary pathways were identified for the binding and dissociation, denoted as “pathway 1” (residues 65-95, including motifs β3, β1’, α1’ and β2’) and “pathway 2” (residues 97-106, α2’ helix) (**Figs. 3 and S2**). These pathways were consistently observed in the binding and dissociation of Nutlin 3, PMI and P53. Binding of Nutlin 3 via pathways 1 and 2 were observed 13 times and 5 times, respectively (**Fig. 3B**). The same pathways 1 and 2 were identified in the simulations of the MDM2-PMI and MDM2-P53 systems. Peptide binding in the MDM2-PMI and MDM2-P53 systems occurred along pathways 1 and 2 for 9 and 7 times, respectively (**Fig. 3B**). Similarly, peptide P53 binding events along pathways 1 and 2 were 8 and 6, respectively (**Fig. 3B**). The same pathways were identified for the dissociation of the MDM2-Nutlin 3, MDM2-PMI and MDM2-P53 systems (**Fig. 3C**). Dissociation of Nutlin-3 via pathways 1 and 2 were observed 11 and 6 times, respectively (**Fig. 3C**). Peptide dissociation in the MDM2-PMI system along pathways 1 and 2 occurred 9 and 9 times, respectively (**Fig. 3C**). Similarly, peptide P53 dissociation along pathways 1 and 2 occurred 6 and 8 times, respectively (**Fig. 3C**).

**Figure 3.**
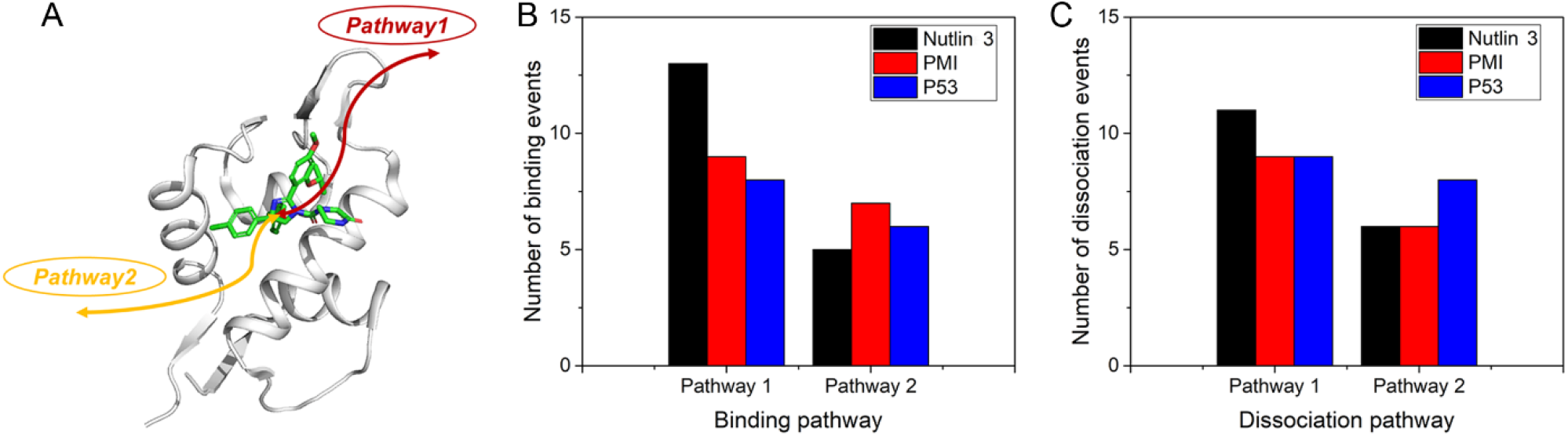
Pathways of ligand/peptide binding and dissociation in the MDM2 protein. (**A**) Cartoon representation of the protein. Binding and dissociation pathways are denoted by the arrow lines. Number of binding (**B**) and dissociation (**C**) events through the different pathways captured by the LiGaMD3 simulations.

### Small-molecule and Peptide binding to the MDM2 involved Induced Fit

After identifying the pathway 1, which involves motifs β3-β1’-α1’-β2’ (residues 65-95), we further investigated the relationship between conformational changes within this region upon small-molecule/peptide binding. Therefore, the ligand RMSD and the number of contacts between the ligand and residues 65-95 in MDM2 (denoted as N_contact_) were used as reaction coordinates to calculate 2D PMF profiles (**Figs. 4A-4C**). Four low-energy states were identified in the 2D PMF profile of the MDM2-Nutlin 3 system including the Bound (“B”), Intermediate (“I1” and “I2”), and Unbound (“U”) (**Fig. 4A**). The ligand RMSD and N_contact_ of these states centered around (3.0 Å, 24), (9.0 Å, 37), (20.1 Å, 0), and (50.0 Å, 0), respectively (**Fig. 4A**). In the MDM2-PMI system, four low-energy states were identified: Bound (“B”), Intermediate (“I1” and “I2”), and Unbound (“U”) (**Fig. 4E**), with the PMI peptide RMSD and N_contact_ centered around (3.5 Å, 60), (9.7 Å, 85), (27.5 Å, 0), and (50 Å, 0), respectively (**Fig. 4E**). Similarly, in the MDM2-P53 system, four low-energy states were observed: Bound (“B”), Intermediate (“I1” and “I2”), and Unbound (“U”) states (**Fig. 4I**), with the P53 peptide RMSD and N_contact_ centered around (4.0 Å, 64), (8.5 Å, 78), (30.5 Å, 0), and (60.8 Å, 0), respectively (**Fig. 4I**).Compared to the Bound state, the intermediate “I1” and “I2” states exhibited significant conformational alterations in the MDM2-Nutlin 3, MDM2-PMI, and MDM2-P53 systems (**Figs. 4B, 4F & 4J**). In the intermediate “I1” and “I2” states, motifs β3-β1’-α1’-β2’ (residues 65-95) in the MDM2-Nutlin3 system moved outward significantly compared to the X-ray Bound structures, resulting in the opening of the binding pocket (**Fig. 4B**).

**Figure 4.**
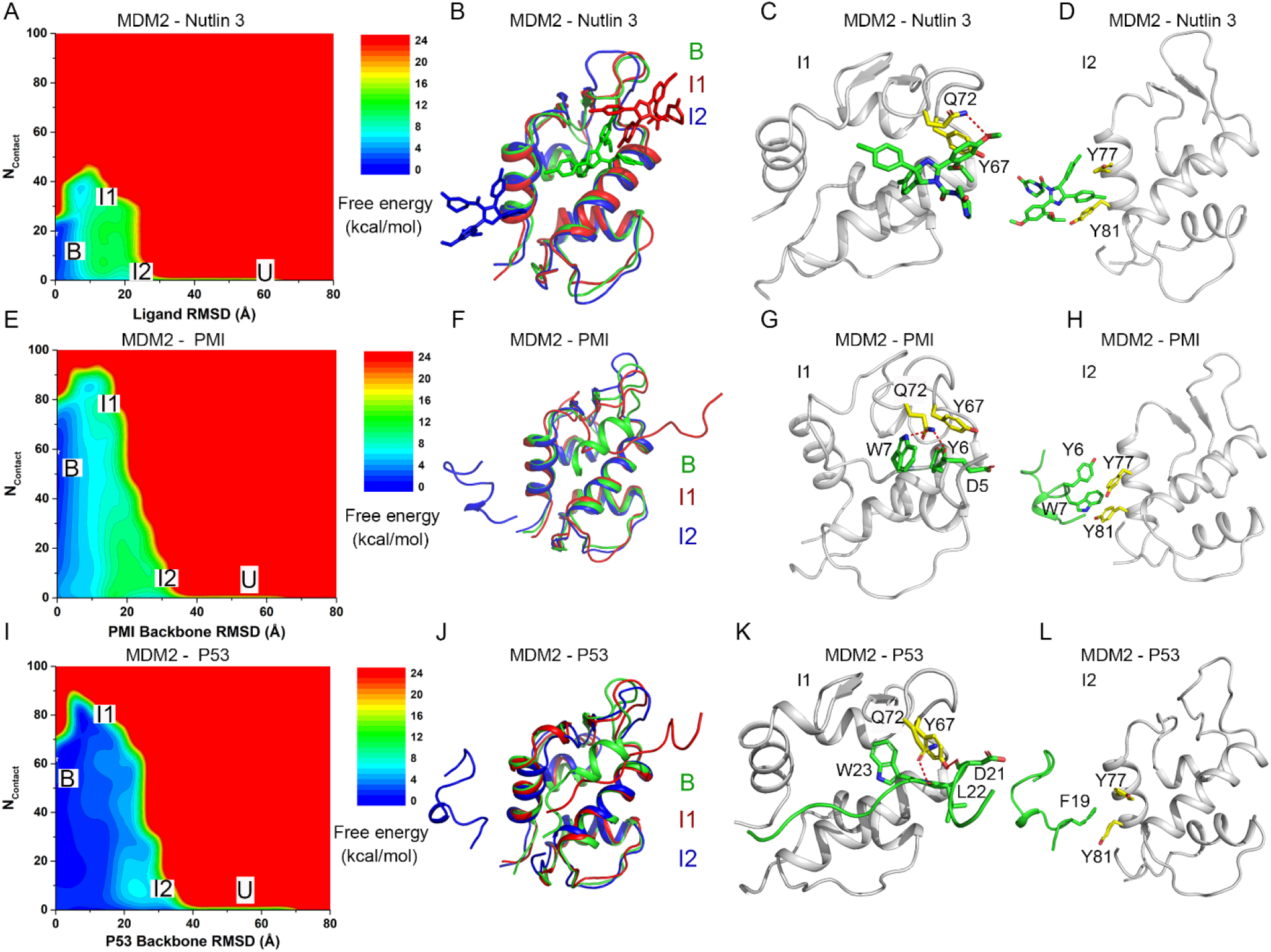
2D Potential of Mean Force (PMF) free energy profiles and low-energy conformational states of ligand/peptide binding to the MDM2: (**A**) 2D PMF profile regarding the ligand heavy atom RMSD and the number of contacts between the ligand and residues 65-95 of MDM2 in the LiGaMD3 simulations of Nutlin 3 binding to the MDM2 protein; (**B**) Low-energy “Intermediate” conformations “I1” (blue) and “I2” (red) as identified from the 2D PMF profiles of Nutlin 3 binding to MDM2 protein; (**C**-**D**) Important ligand-MDM2 interactions in the low-energy conformations “I1” (**C**) and I2 (**D**). (**E**) 2D PMF profile regarding the peptide backbone RMSD and the number of contacts between the PMI and residues 65-95 of the MDM2 in the LiGaMD3 simulations of PMI binding to the MDM2 protein; (**F**) Low-energy “Intermediate” conformations “I1” (blue) and “I2” (red) as identified from the 2D PMF profiles of PMI binding to MDM2 protein; (**G**-**H**) Important PMI-MDM2 interactions in the low-energy conformations “I1” (**G**) and I2 (**H**). (**I**) 2D PMF profile regarding the peptide backbone RMSD and the number of contacts between the P53 and residues 65-95 of the MDM2 in the LiGaMD3 simulations of P53 binding to the MDM2 protein; (**J**) Low-energy “Intermediate” conformations “I1” (blue) and “I2” (red) as identified from the 2D PMF profiles of P53 binding to MDM2 protein; (**K**-**L**) Important P53-MDM2 interactions in the low-energy conformations “I1” (**K**) and I2 (**L**).

Two critical interactions were identified in “I1” state: a hydrogen bond interaction between MDM2:Q72 and Nutlin 3 and aromatic interaction between MDM2:Y67 and Nutlin 3 (**Fig. 4C**). While in the “I2” state, Π-Π interactions were observed between MDM2:Y77 or MDM2:Y81 and Nutlin 3 (**Figs. 4C&4D**). In the MDM2-PMI system, significant conformational changes occurred upon peptide PMI binding, involving particularly motifs β3-β1’-α1’-β2’ (residues 65-95), resulting in distinct open and closed conformations in the “I1” and “I2” states, respectively (**Fig. 4F**). In the “I1” state, hydrogen bonds were formed between MDM2:Q72 and PMI:W7, MDM2:Q72 and PMI:D5 (**Figs. 4G**). In the “I2” state, Π-Π interactions were formed between MDM2:Y77-PMI:Y6, MDM2:Y81 and PMI: W7 (**Figs. 4H**). In the MDM2-P53 system, significant conformational changes occurred upon peptide P53 binding, involving particularly motifs β3-β1’-α1’-β2’ (residues 65-95), resulting in a more closed conformation in the “I1” and “I2” states (**Fig. 4J**). In the “I1” state, hydrogen bonds were formed between MDM2:Q72 and P53:L22, MDM2:Y67 and P53:D21 (Fig. 4K). In the “I2” state, Π-Π interactions were formed between MDM2:Y77 and P53:F19, MDM2:Y81 and P53:F18 (**Figs. 4K&4L**). Hence, residues Q72, Y67, Y77 and Y81 in the MDM2 protein play pivotal roles during ligand binding in the intermediate conformational states.

In order to further explore the mechanism of ligand binding to the MDM2, we computed 2D PMF free energy profiles to characterize conformational changes of both the protein and ligand during binding. The intermediate “I1” and “I2” states showed quite large conformational changes in the motifs β3-β1’-α1’-β2’ (residues 65-95). Therefore, we calculated 2D PMF profiles regarding the RMSD of the ligand and the MDM2 motifs β3-β1’-α1’-β2’ (residues 65-95) RMSD (denoted as Loop RMSD) relative to the experimental bound structures with the protein aligned (**Figs. 5A-5C**). For the MDM2-Nutlin 3 system, three low-energy states were identified from the 2D PMF profile, including the Bound, Intermediate “I2” and Unbound (**Figs. 5A**). The protein motifs β3-β1’-α1’-β2’ at the peptide-binding site adopted the “Open” conformation in the I2 state (**Figs. 5A** and **5B**). The peptide and loop RMSDs centered around (4.5 Å, 1.0 Å), (18.0 Å, 2.9 Å) and (59.0 Å, 2.0 Å) in the Bound “B”, Intermediate “I2” and Unbound “U” states, respectively (**Fig. 5A**). For the MDM2-PMI system, three low-energy states were identified from the 2D PMF profile, including the Bound “B”, Intermediate “I1” and Unbound “U”. The peptide and loop RMSDs centered around (5.0 Å, 0.8 Å), (10.2 Å, 3.0 Å) and (58.2 Å, 2.0 Å) in the “B”, “I1” and “U” states, respectively (**Figs. 5B**). Four low-energy conformational states were identified in the MDM2-P53 system. The peptide and loop RMSDs centered around (5.2 Å, 0.8 Å), (10.5 Å, 3.9 Å), (31.0 Å, 4.0 Å) and (61.0 Å, 1.9 Å) in the Bound “B”, Intermediate “I1” and “I2”, and Unbound “U” states, respectively (**Fig. 5C**).

**Figure 5.**
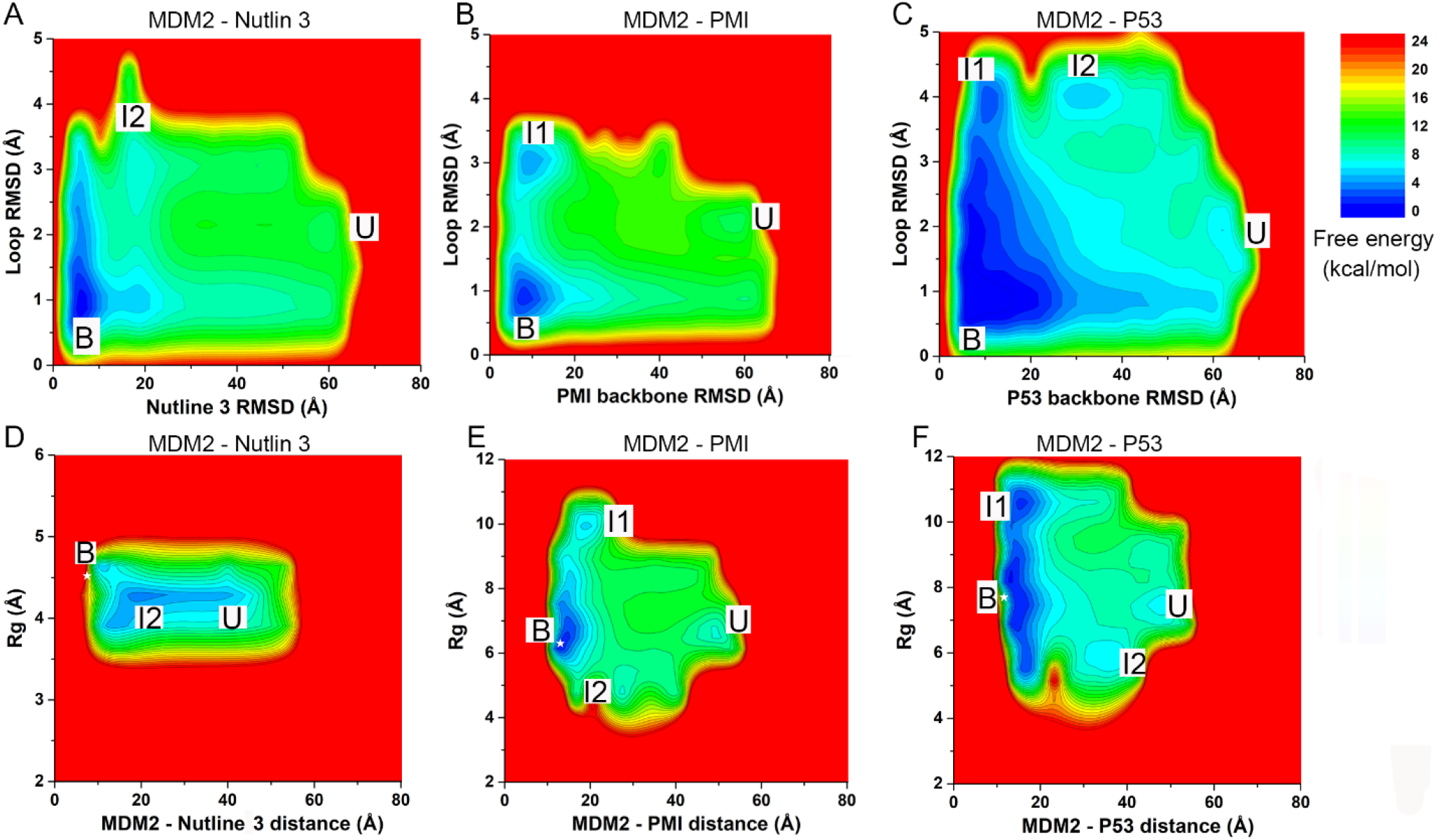
(**A-C**) 2D PMF profiles regarding the ligand heavy atom RMSD or peptide backbone RMSD and the MDM2 loop (residues 65-95) RMSD relative to their corresponding experimental bound structure in the LiGaMD3 simulations of Nutlin 3(**A**), PMI peptide (**B**) and P53 peptide (**C**) binding to the MDM2 protein; (**D-F**) 2D PMF profiles regarding the distance between the ligand/peptide and MDM2 binding pocket and the *Rg* of the substrates in the LiGaMD3 simulations of Nutlin 3 (**D**), PMI peptide (**E**) and P53 (**F**) binding to the MDM2 protein.

In addition, we examined the conformational dynamics exhibited by the small molecule and peptides during their binding processes. In this regard, the ligand radius of gyration (*R*_*g*_) was calculated and monitored for possible conformational changes. The small-molecule/peptide *R*_*g*_ and the center-of-mass distance between protein pocket and ligand (denoted as d_MDM2-substrate_) were used as reaction coordinates to calculate 2D PMF profiles. From the reweighted 2D PMF profiles (**Fig. 5D-5F**), we identified a low-energy “Bound” state in all three systems, for which the d_MDM2-substrate_ and *R*_*g*_ in the MDM2-Nutlin3, MDM2-PMI and MDM2-P53 systems centered around (8.6 Å, 4.4 Å), (14.0 Å, 6.3 Å) and (13.0 Å, 8.3 Å), respectively. This suggested that successful sampling of complete small-molecule/peptide binding was captured in the LiGaMD3 simulations. Notably, in the intermediate states, the peptides sampled a wider range of *R*_g_ and the protein motifs β3-β1’-α1’-β2’ exhibited higher RMSDs compared with the bound states. Therefore, the small-molecule and peptides binding to the MDM2 protein showed predominantly an “induced-fit” mechanism.

## Discussions

We have presented a new LiGaMD3 method to improve sampling efficiency and accurately predict thermodynamic and kinetic properties associated with the binding of small molecules and highly flexible peptides. LiGaMD3 works by selectively boosting the essential non-bonded interaction potential energy of the ligand, as well as the remaining non-bonded potential energy and all the bonded potential of the system. Non-bonded potential interactions play a critical role in ligand dissociation and rebinding, while the bonded potentials mainly contribute to conformational changes of the system. Utilizing microsecond timescale simulations, LiGaMD3 effectively captures repetitive dissociation and rebinding processes of both small molecules and peptides in three model systems of MDM2 bound by different small molecules and flexible peptides. These simulations then enable accurate predictions of ligand/peptide binding free energies and kinetic rate constants.

LiGaMD3 simulations revealed the critical role of nonbonded potentials in governing ligand dissociation and rebinding process, being consistent with previous computational findings^23, 32, 62^. Non-bonded interactions have been recognized as one of the main factors that govern the ligand binding to its target protein^43^. Furthermore, our simulations identified multiple pathways for ligand binding and dissociation and revealed an “induced-fit” mechanism of ligand binding, being consistent with earlier simulation results^15, 33, 63^. Compared with the cMD^64^, Metadynamics^65^, Weighted Ensemble,^66^ MSM^30^ and Replica Exchange MD simulations^41^, LiGaMD3 offers a more efficient and user-friendly approach. LiGaMD3 also shows advantages over previous LiGaMD, particularly in its ability to accurately capture peptide binding to proteins. While microsecond cMD simulations have proven effective in capturing small molecule and highly flexible peptide binding to target proteins, the slower kinetics of ligand dissociation remain beyond the accessible timescale of cMD. Weighted Ensemble^32^ and MSM^30^ methods have shown promise in accurately predicting small molecule and peptide binding kinetics, but typically require extensive computational resources, often involving tens-of-microsecond simulations^30^. Metadynamics, with carefully designed CVs, can efficiently capture both ligand binding and unbinding. However, the predefined CVs may impose constraints on binding pathways and conformational space. The approach may also encounter challenges such as the “hidden energy barrier” problem and slow convergence if important CVs are omitted.^67^ Overall, previous methods have been computationally demanding, necessitating significantly longer simulations to adequately characterize ligand binding thermodynamics and kinetics. LiGaMD3 captures the repetitive small-molecule and peptide dissociation and binding events within only microsecond simulations, offering an efficient approach to characterizing ligand binding dynamics and extending the capabilities of the LiGaMD methodology to binding of highly flexible peptides.

## Acknowledgements

This work used supercomputing resources with allocation award TG-MCB180049 and BIO210039 through the Advanced Cyberinfrastructure Coordination Ecosystem: Services & Support (ACCESS) program, which is supported by National Science Foundation grants #2138259, #2138286, #2138307, #2137603, and #2138296, and project M2874 through the National Energy Research Scientific Computing Center (NERSC), which is a U.S. Department of Energy Office of Science User Facility operated under Contract No. DE-AC02-05CH11231, and the Research Computing at the UNC-Chapel Hill. This work was supported in part by the National Institutes of Health (R01GM132572) and National Science Foundation (2121063).

## Supporting Information

Example simulation files of LiGaMD3 simulations of the MDM2-Nutlin 3 system, **Tables S1, S2, Figures S1 and S2** are provided in the Supporting Information. This information is available free of charge via the Internet at http://pubs.acs.org.

## TOC

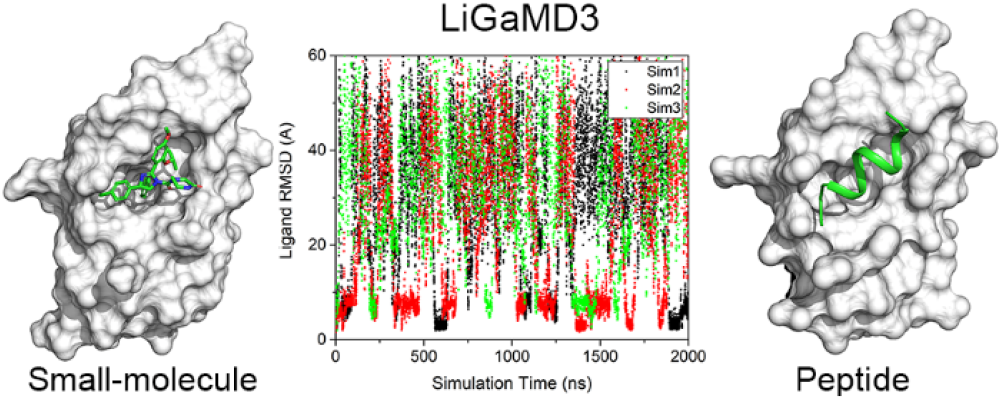

