## Supporting information for "Ligand Gaussian accelerated Molecular Dynamics 3 (LiGaMD3): Improved Calculations of Binding Thermodynamics and Kinetics of Both Small Molecules and Flexible Peptides"

**Table S1** The ligand bound and unbound time periods ($\tau_B$ and $\tau_U$) recorded from LiGaMD3 simulations of the ligand/peptide-MDM2 binding systems.

| System | Method | ID | $\tau_B$ (ns) | $\tau_U$ (ns) |
|---|---|---|---|---|
| MDM2-Nutlin | LiGaMD | Sim1 | 74,321,19,172,146 | 42.4,395,56,97,36,632 |
| | | Sim2 | 187,1026,209 | 87,317,184 |
| | | Sim3 | 397,334,706 | 26,66,67,404 |
| MDM2-Nutlin | LiGaMD3 | Sim1 | 14,55.7,23,83,23,115 | 25,240.9,196,600,623 |
| | | Sim2 | 61.6,53,87,12,65,47,8 | 272,137,338,20,219,137,164 |
| | | Sim3 | 14,30,30,24,23 | 189,628,551,119,392 |
| MDM2-PMI | LiGaMD3 | Sim1 | 360.7,191.6,64.29,10.5,32.9,17.1 | 85.2,17.1,26.3,40.5,43.3,101.5 |
| | | Sim2 | 456.4,47.2,18.4 | 261.1,110.6,96.3 |
| | | Sim3 | 68.17,147.0,57.6,10.5,9.8,45.0,8.9 | 129.8,35.4,56.6,45.4,108.2,59.2,221.0 |
| MDM2-P53 | LiGaMD3 | Sim1 | 60,17,29,25,30 | 250,136,249,153.7,534,380 |
| | | Sim2 | 379,41,30,17 | 306,201,390,197,434 |
| | | Sim3 | 38.5,23.7,88,24,84 | 100.7,115,118,207,407,787 |

**Table S2** Energy barriers of ligand/peptide-MDM3 dissociation ("off") and binding ("on") calculated from the reweighed (Δ***F***) and modified (no reweighting, Δ***F****) free energy profiles, curvatures of the reweighed ($w$) and modified ($w^*$) free energy profiles near the ligand/peptide Bound ("B"), Barrier ("Br") and Unbound ("U") states, and the ratio of apparent diffusion coefficients calculated from the LiGaMD3 simulations without reweighting (modified, $D^*$) and with reweighting ($D$).

| Sim | | Δ***F*** (kcal/mol) | | Δ***F**** (kcal/mol) | | ***w*** | | | ***w**** | | | ***D*/D*** | |
|---|---|---|---|---|---|---|---|---|---|---|---|---|---|
| | | **Off** | **On** | **Off** | **On** | **B** | **Br** | **U** | **B** | **Br** | **U** | **Off** | **On** |
| MDM2-Nutlin | LiGaMD | 9.10± 1.26 | 2.77 ±1.76 | 2.68 ±0.25 | 0.63 ±0.16 | 1.54 ±0.36 | 0.30 ±0.30 | 0.22 ±0.17 | 0.78 ±0.028 | 0.069 ±0.009 | 0.020 ±0.005 | 1.43 ±0.44 | 0.16 ±0.03 |
| | LiGaMD3 | 9.65 ±0.76 | 0.79 ±0.10 | 0.76 ±0.18 | 0.33 ±0.15 | 10.28 ± 0.60 | 0.12 ±0.07 | 0.012 ±0.006 | 1.16 ±0.27 | 0.14 ±0.05 | 0.032 ±0.009 | 0.96 ±0.81 | 0.18 ±0.05 |
| MDM2-PMI | LiGaMD3 | 9.05± 0.23 | 0.89 ±0.07 | 1.38 ±0.12 | 0.23 ±0.06 | 0.94±0 .53 | 0.0097 ±0.0043 | 0.048 ±0.020 | 7.21 ±0.30 | 0.066 ±0.010 | 0.025 ±0.003 | 0.13 ±0.03 | 0.17 ±0.004 |
| MDM2-P53 | LiGaMD3 | 7.23± 0.40 | 0.67 ±0.18 | 1.16 ±0.04 | 0.093 ±0.031 | 0.43 ±0.20 | 0.023 ±0.012 | 0.023 ±0.018 | 7.04 ±0.10 | 0.062 ±0.018 | 0.026 ±0.002 | 0.25 ±0.08 | 0.23 ±0.20 |

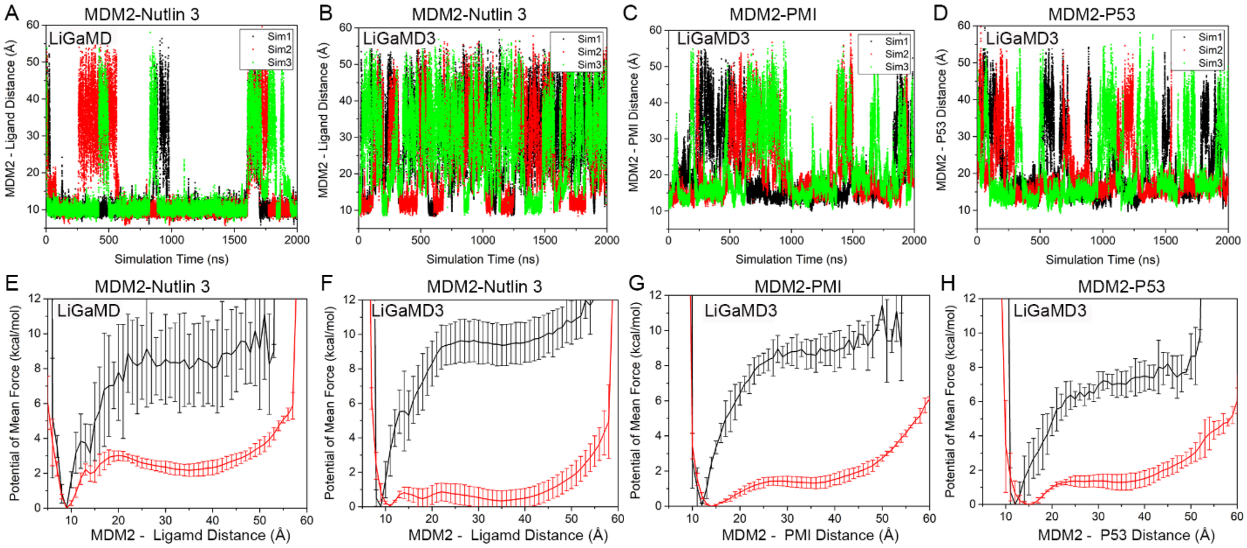


**Figure S1.** LiGaMD3 simulations captured repetitive dissociation and binding of the Nutlin 3 small molecule and highly flexible PMI and P53 peptides to the MDM2 protein: (**A**-**B**) time courses of the distance between the MDM2 and small molecule Nutlin 3 calculated from three independent 2 μs LiGaMD (**A**) and LiGaMD3 (**B)** simulations; (**C**-**D**) time courses of the distance between the MDM2 and peptide from three independent 2 μs LiGaMD3 simulations of (**C**) PMI and (**D**) P53 binding to MDM2; (**E**-**F**) The corresponding reweighted (black) and non-reweighted (red) PMF profiles of the distance between MDM2 and ligand averaged over three LiGaMD (**E**) and LiGaMD3 (**F**) simulations of Nutlin 3 binding to MDM2; (**G**-**H**) )The corresponding PMF profiles of the MDM2-peptide distances averaged over three LiGaMD3 simulations of (**G**) PMI and (**H**) P53 binding to MDM2. Error bars are standard deviations of the free energy values calculated from three LiGaMD and LiGaMD3 simulations.

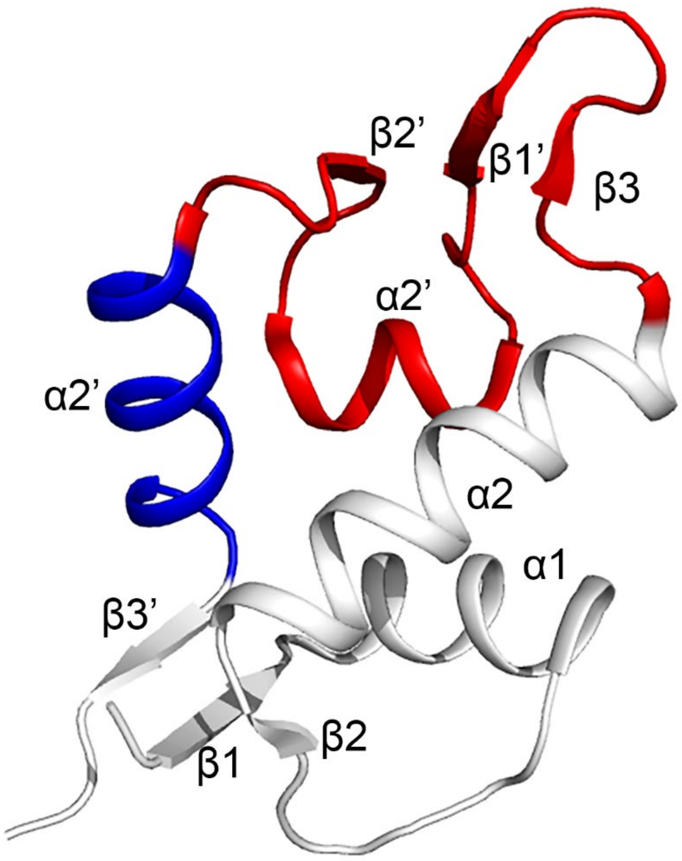


**Figure S2.** Cartoon representation of the MDM2 with each motif labeled. Residues predominantly involved in "pathway 1" were colored in red, including the β3, β1' and β2' strands and the α1' helix (residues 65-95). Residues mainly involved in "pathway 2" were colored in blue, including the α2' helix (residues 97-106).
